# An interpretable machine learning framework for dog breed inference and ancestry decomposition

**DOI:** 10.64898/2026.06.03.729926

**Authors:** Yiming Bian, Rob Bierman, Noah Snyder-Mackler, Daniel Promislow, Elinor Karlsson, Dog Aging Project Consortium, Joshua M. Akey

**Author notes:** Dog Aging Project Consortium collaborators and affiliations are listed at the end of this paper.

## Abstract

The over 300 currently recognized breeds of domesticated dogs are the culmination of centuries of intense artificial selection and recurrent population bottlenecks. While breed labels are widely used in genetic and veterinary studies, inferring breed identity from genomic data remains challenging due to the high dimensionality of genotype data, uneven sampling across breeds, and admixture resulting in mixed-breed individuals. Here, we present an interpretable machine learning framework to infer dog breed labels from genome-wide SNP data. Our approach combines dimensionality reduction with a multi-output random forest model that maps genetic variation to a continuous representation of breed membership, enabling both classification and mixed-breed inference. We apply this framework to the Dog Aging Project (DAP) dataset of 6,572 purebred and mixed-breed dogs across 100 breed classes, achieving 91.7% accuracy with an overlap-based metric, outperforming an ADMIXTURE-based benchmark that achieved 87.8% accuracy. Notably, we find that as few as 150 informative SNPs are sufficient to achieve near-maximal predictive performance, highlighting the highly structured nature of canine genetic variation. We also introduce a SNP importance score metric that links model predictions back to individual genetic variants. Analysis of top-ranked variants reveals loci previously associated with morphological, pigmentation, and behavioral traits, as well as candidate loci lacking prior phenotypic annotation, supporting both the biological relevance and discovery potential of the framework. Together, these results demonstrate that our framework provides an accurate, flexible, and interpretable approach to predict breed ancestry, with applications in veterinary genomics, canine population genetics, and the identification of loci underlying hallmark breed phenotypes.

## Introduction

Domestic dogs (*Canis familiaris*) are among the most genetically and phenotypically diverse mammalian species, owing to domestication, intense artificial selection, and repeated population bottlenecks^1,2^. This history has produced more than 300 recognized breeds, each shaped by distinct evolutionary and demographic processes and characterized by unique genetic profiles^3^. Accurate breed prediction, therefore, has practical value for veterinary research and preventive care^4,5^, while also providing an important framework for studying the genetic architecture of breed-associated traits^6^. Although pedigree records and morphological characteristics can be informative, they are often unreliable for mixed-breed dogs or individuals with uncertain ancestry^7^. Genomic data, particularly single nucleotide polymorphisms (SNPs), provide a more objective and reproducible basis for breed prediction.

Despite its potential power, SNP-based breed prediction remains challenging. A major obstacle is the extreme dimensionality of canine genomic data, in which the number of SNPs far exceeds the number of available samples. This imbalance complicates model training, increases the risk of overfitting, and imposes substantial computational and memory demands. A second challenge concerns the quality and distribution of breed labels. Many datasets rely on owner-reported breed identities^6,8^, which may be inaccurate, especially for mixed-breed or visually ambiguous dogs. In addition, breed representation is highly uneven, with popular breeds such as Labrador Retrievers, German Shepherds, and Golden Retrievers often well sampled, whereas rarer breeds, such as Maremmas, Barbets, and Komondors, may be represented by only a few individuals^9^ or not sampled at all. Collectively, these issues limit statistical power for underrepresented groups and complicate prediction in mixed-breed dogs, highlighting the need for dimensionality reduction, feature selection, and robust predictive models tailored to structured genomic data^10^.

A range of approaches have been used to infer the ancestry of individual dogs from genome-scale data^1,3,6,11–13^ (typically SNPs, although other classes of variants have also been used^1^).

Classic population-genetic approaches such as STRUCTURE^14^, ADMIXTURE^15^, and fastSTRUCTURE^16^ were developed to infer ancestry and admixture proportions from genome-wide genotype data and have subsequently been used to infer breed labels in dogs and to characterize canine population structure^3,6^. More recent studies have explored supervised and machine-learning approaches for assigning breed labels or estimating breed composition from SNP data^12,17^. However, many existing methods either emphasize single-label prediction, which makes them less suitable for mixed-breed dogs, or provide limited interpretability at the variant level. Indeed, there is growing interest in interpretable machine-learning frameworks^18^ that not only achieve accurate predictions but also identify the specific genetic variants that contribute most strongly to model performance. Thus, these observations motivate the need for additional methodological development that combines ancestry-aware prediction, computational scalability, biological interpretability, and robustness to reduced-marker representations. Such approaches are particularly relevant in dogs, where strong breed structure and extensive linkage disequilibrium (LD)^10^ suggest that a relatively small number of highly informative variants may capture much of the predictive signal encoded across the genome.

Here, we present an interpretable machine-learning framework that uses genome-wide SNP data to predict breed labels for individual dogs. We apply our approach to 6,572 purebred and mixed-breed dogs with low-coverage whole-genome sequencing (WGS) data from the Dog Aging Project (DAP)^9^, and demonstrate that it enables robust breed prediction, outperforming current state-of-the-art approaches^13^. We also develop a SNP importance score that identifies genetic variants informative for breed prediction, detecting loci previously associated with hallmark breed phenotypes and genes whose phenotypic consequences are unknown. We further demonstrate that near-maximal predictive performance can be achieved using only a few hundred informative SNPs, indicating that much of the genomic signal underlying breed identity is concentrated in a relatively small set of highly informative variants. Together, these results establish a compact and interpretable framework for breed inference and provide insight into the genetic architecture of domesticated dog populations.

## Results

### Overview of the breed prediction framework

To infer breed identity from genomic data, we developed an interpretable machine-learning framework that maps genome-wide SNP variation onto a continuous representation of breed membership (Figure 1a). The model is trained on a panel of purebred dogs, where each breed represents a distinct class. We also include an "unknown" class during training to accommodate cases where a dog contains ancestry from a breed not represented in the training panel (Figure 1a). Rather than assigning a single discrete class label, the model produces a vector of breed-membership scores that can represent both purebred and mixed-breed dogs through downstream post-processing.

**Figure 1.**
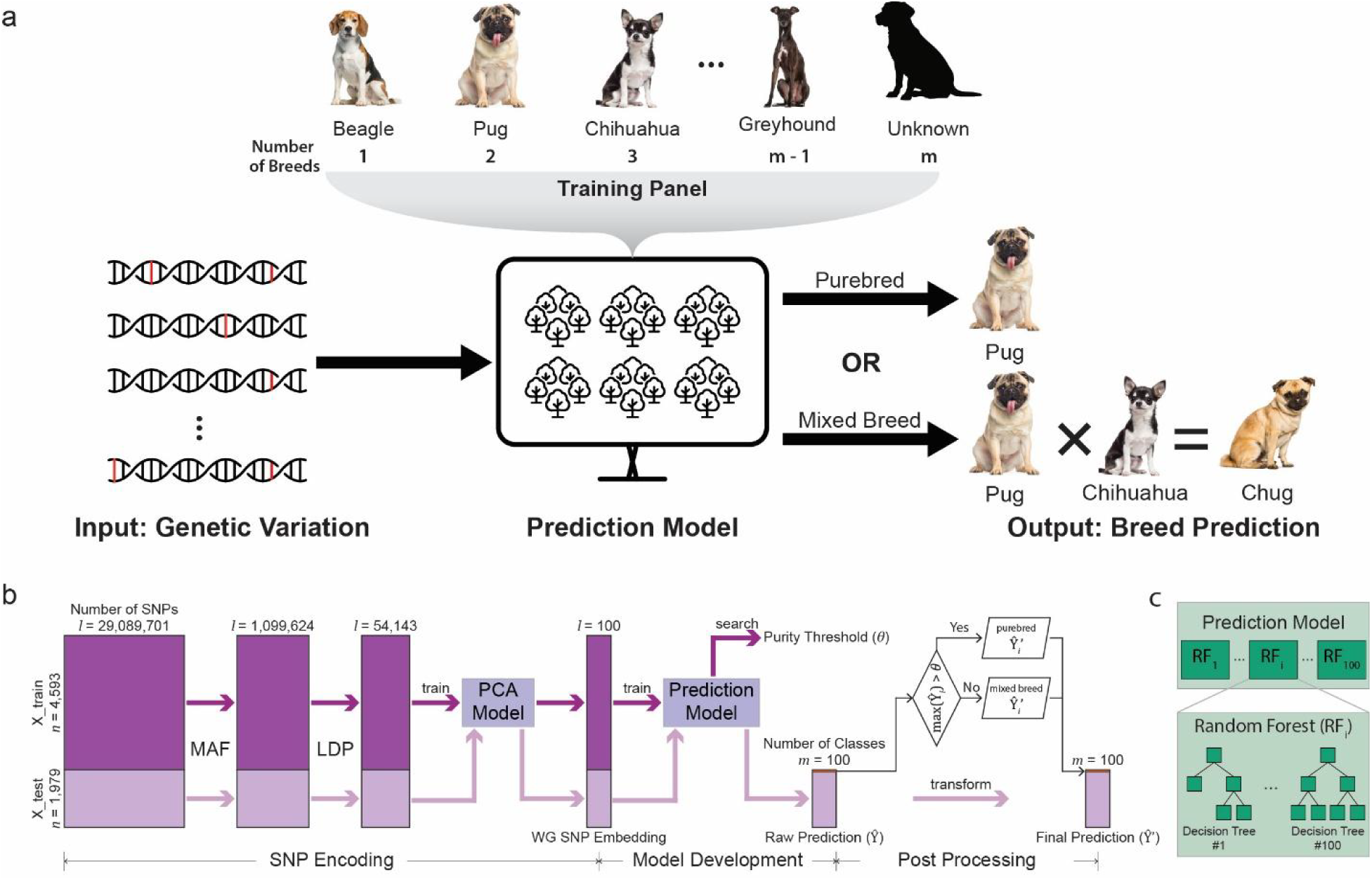
Breed-prediction framework. (a) Schematic of dog breed prediction from genome-wide SNP data. The model is trained on a reference panel of m breed classes, including m − 1 recognized breeds and one “Unknown” class, and outputs either a purebred or a mixed-breed prediction. (b) Overall workflow for SNP encoding, model development and post-processing. (c) Architecture of the prediction model, comprising m breed-specific random forests. In our main model, m = 100 and each random forest consists of 100 decision trees.

A more detailed schematic of the prediction model framework is shown in Figure 1b, which consists of three primary steps: 1) SNP encoding, 2) model development, and 3) post-processing to obtain the final prediction. To reduce the dimensionality of the SNP data and remove variants with redundant information, we filter SNPs using a minor allele frequency (MAF) threshold and then perform LD pruning. The resulting SNP matrix is transformed into a compact whole-genome SNP embedding via principal component analysis (PCA), yielding a reduced-dimensional representation of genome-wide genetic variation for downstream model training.

This embedding was used as input to a collection of breed-specific random forests^19^, which generate continuous breed-membership scores for each individual. Random forests are ensemble machine-learning models that combine predictions from many decision trees, allowing complex and nonlinear relationships between genetic variants and breed identity to be learned while reducing overfitting and improving predictive robustness.

Figure 1c illustrates the ensemble architecture of the prediction framework, in which each breed-specific model comprises a random forest of multiple decision trees trained on the reduced SNP representation. A post-processing procedure is subsequently applied to transform these raw predictions into final breed assignments for both purebred and mixed-breed dogs. Together, this workflow enables scalable breed prediction from high-dimensional genomic data while retaining a direct connection between model predictions and informative genetic variants. Note that Figure 1b presents the dimensionality and other characteristics of the empirical data at each step of the analysis, as described in more detail below. Below, we describe the SNP encoding, model development, and post-processing analyses in more detail.

### Encoding high-dimensional SNP data into a compact representation

The large number of SNPs across the canine genome poses a problem for supervised prediction models. For example, the Dog Aging Project (DAP) dataset^9,20^ includes 7,627 dogs with DNA sequencing data and more than 29 million SNPs across the genome. Thus, it is necessary to reduce this feature space, which we accomplished by transforming the raw genotype matrix into a compact SNP embedding that captures the major axes of genetic variation while remaining computationally tractable. Specifically, we applied three successive filtering and compression steps: minor allele frequency (MAF) thresholding, LD pruning (LDP) performed separately on each chromosome, and PCA on the concatenated post-pruning SNP set^21^ (Figure 1b).

We evaluated a range of MAFs, LDP thresholds, and PCA dimensions (Table 1). Unless otherwise stated, we used a MAF ≥ 0.40, LDP threshold of 𝑟^2^ ≥ 0.10 (see Methods), and 100 PCs (Table 1) for SNP encoding. When choosing these values, we considered the model’s predictive performance, robustness, and size (with a preference for smaller, more compact models). These factors and alternative parameter settings are described in the Discussion.

**Table 1.**
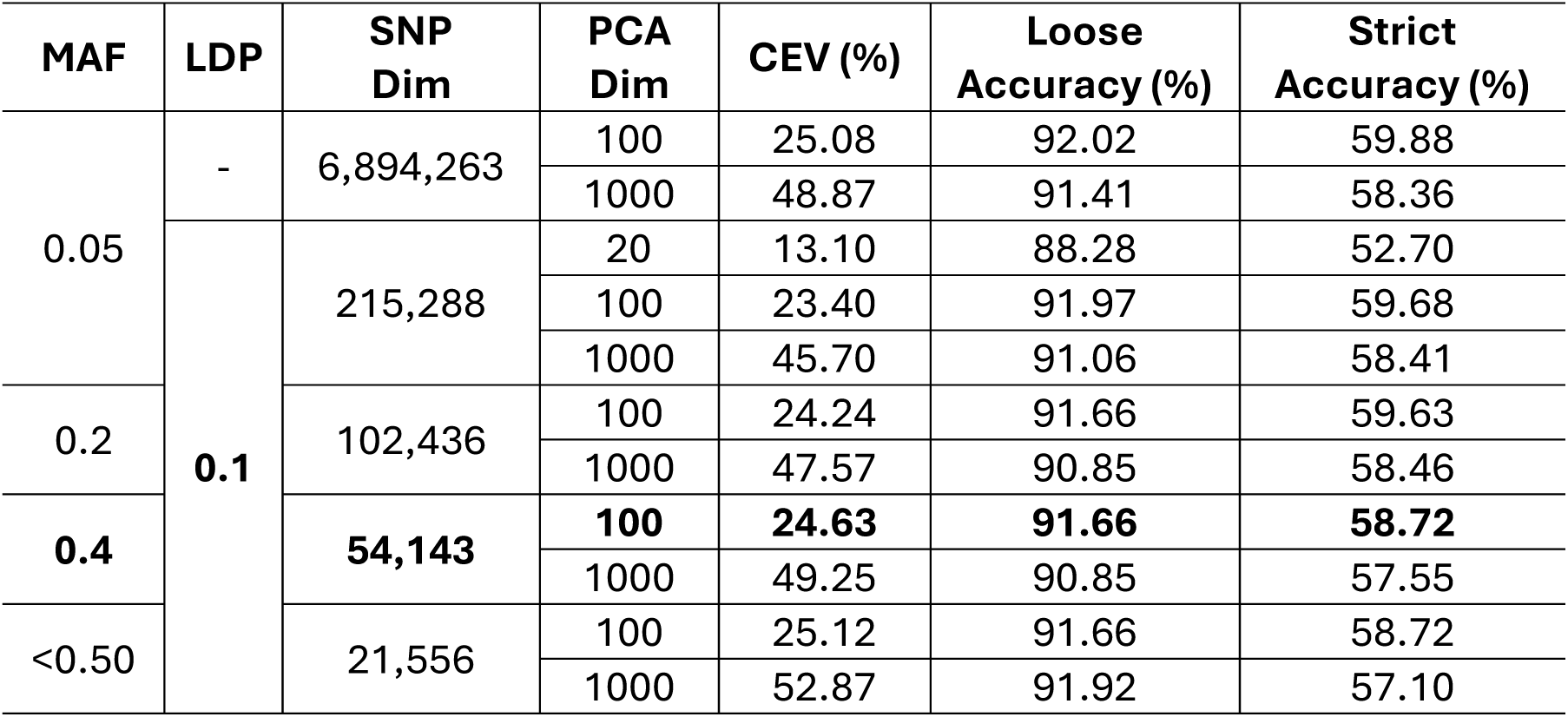
Comparison of SNP encoding parameter settings and their effect on model performance. Shown are minor allele frequency (MAF) thresholds, linkage disequilibrium pruning (LDP) threshold, the resulting SNP dimension, PCA dimension, cumulative explained variance (CEV), and prediction performance measured by loose and strict accuracy. The bolded row indicates the final encoding configuration selected as a compromise between predictive performance and representational compactness.

Although 0.40 is a high MAF threshold, it is important to note that this is the global minor allele frequency across all DAP dogs. Because breeds exhibit significant population structure^1–3^, this global MAF threshold enriches for variants with large allele-frequency differences across breeds. For example, consider a biallelic SNP with alleles “1” and “2” and corresponding allele frequencies of *p*;_1_and *p*;_2_, respectively. In the extreme case where half of the breeds are fixed for allele 1 (*p*;_1_ = 1, *p*;_2_ = 0) and the other half are fixed for allele 2 (*p*;_1_ = 0, *p*;_2_ = 1 ), the global MAF is (1 + 0) / 2 = 0.50. After MAF and LDP filtering, we reduced the genome-wide variant set from more than 29 million to 54,143 SNPs, which served as input for PCA (Figure 1b). The resulting 100-dimensional representation explained 24.6% of the variance (Table 1) in the filtered SNP matrix and was used as the input for all downstream breed-prediction analyses. To visualize the population genetic structure within and between breeds contained within the filtered set of 54,143 SNPs, we constructed interactive three-dimensional scatter plots of the top three principal components (see supplementary data).

### Dataset preparation

Among the 7,627 dogs in the DAP dataset, 3,985 were reported as purebred, spanning 233 breeds, and 3,642 were reported as mixed-breed, encompassing 1,273 mixed-breed combinations. The mixed-breed labels comprise combinations of 186 pure breeds, 165 of which overlap with the 233 pure breeds represented among the purebred dogs. However, many breed classes were represented by only a few samples. For example, 85 of the 233 pure breeds and 990 of the 1,273 mixed-breed classes were represented by fewer than three dogs. Given this substantial class sparsity, we restricted model development to a large and relatively well-represented subset of the full dataset. Specifically, we defined 100 purebred classes (m = 100), consisting of the 99 pure breeds with the largest sample sizes together with an additional “Unknown” class (Figure 1a and 1b). We then retained all mixed-breed samples whose reported parental breeds were drawn from these 100 classes. The number of samples in each class is provided in Table S1. This procedure yielded a final dataset of 6,572 dogs, which was partitioned into a training set of 4,593 samples (70%) and a testing set of 1,979 samples (30%), as shown in Figure 1b. The partition was randomized subject to the constraint that every purebred class appeared in both sets. Mixed-breed classes represented by a single sample were assigned randomly to one set or the other. In both subsets, the overall composition was similar, with approximately 55% purebred and 45% mixed-breed dogs.

To encode owner-reported breed labels, we represented each sample as a 100-dimensional vector. Purebred dogs were assigned one-hot labels, with the entry corresponding to the reported breed set to 1 and all others set to 0. Mixed-breed dogs were encoded by assigning a value of 0.5 to each reported parental breed and 0 to all other entries. Here, the value 0.5 is used only to represent a weaker breed-specific signal relative to the value 1 assigned to purebred labels. It should not be interpreted as an estimate of true admixture proportion. Figure 2a illustrates label construction for purebred and mixed-breed dogs.

**Figure 2.**
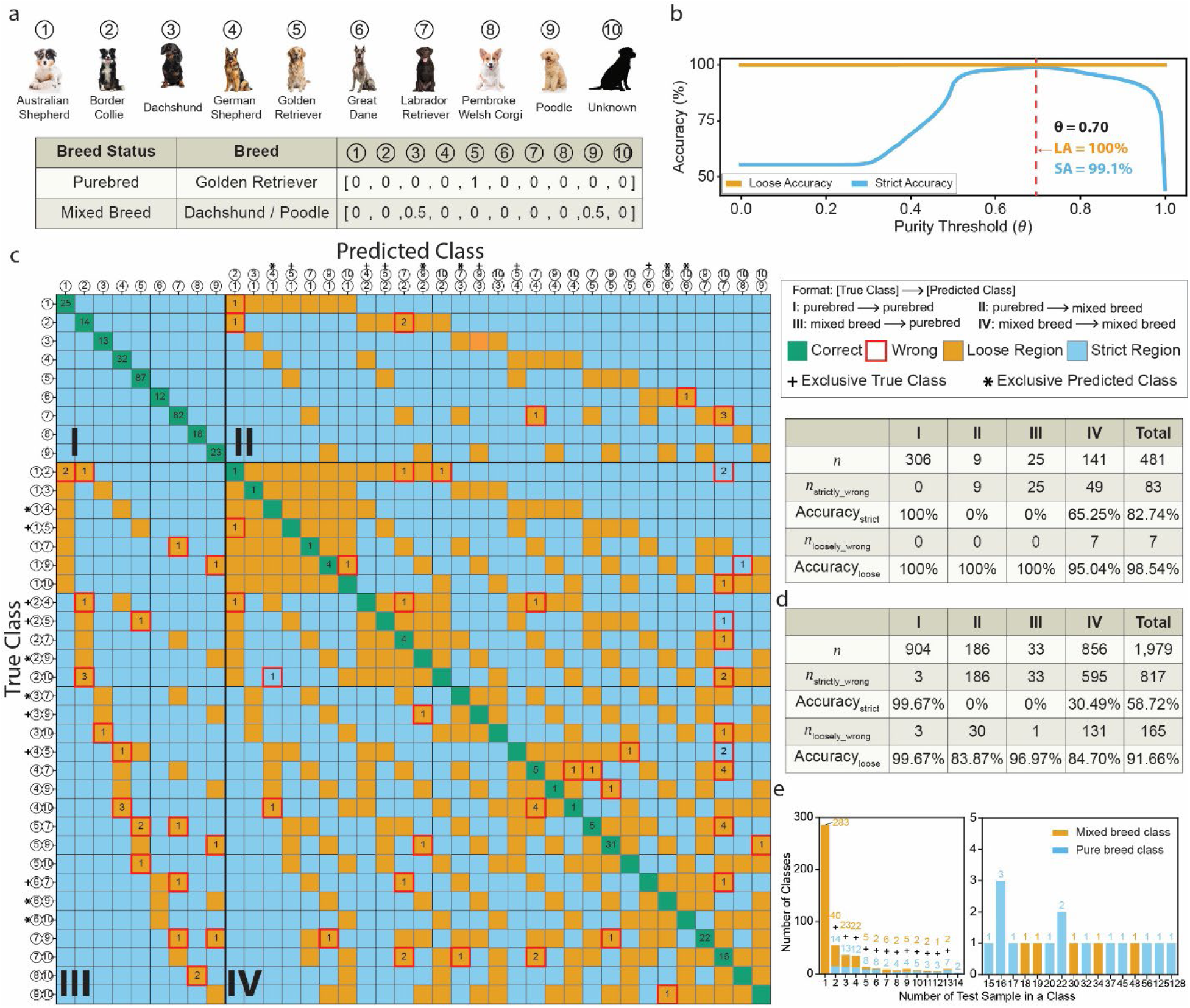
Label construction, threshold selection, and model evaluation. (a) An example of how purebred and mixed-breed labels are constructed (*m* = 10). (b) Purity-threshold search based on loose and strict accuracy. (c) Prediction map and performance for a subset of breeds (*m* = 10). The two axes correspond to the true class labels and predicted class labels, respectively. Labels present only among the true classes are marked with a plus sign (+), whereas labels present only among the predicted classes are marked with an asterisk (*). Each cell is color-coded according to prediction status, and incorrect predictions, including both loosely and strictly incorrect ones, are further highlighted by red boxes. (d) Section-wise performance of the full model for the complete m = 100 prediction task. (e) Distribution of breed classes in the test set; the right panel enlarges classes with at least 15 test samples.

### Breed prediction model

For each dog, the model input is a whole-genome SNP embedding vector of length 100, and the target output is a breed label vector of length 100. As shown in Figure 1b, model development comprises both training and inference. The prediction framework itself consists of 100 independent random forest regressors, one for each breed class, with each regressor containing 100 decision trees (Figure 1c). For a given sample, the 𝑖^𝑡ℎ^ random forest produces a scalar output, denoted *ŷ_i_* (0 ≤ *ŷ_i_* ≤ 1), which reflects the strength of evidence that the sample contains ancestry from the corresponding breed signal. Collecting these outputs across all classes yields the raw prediction vector *Ŷ* = [*ŷ*_1_, … , *ŷ*_100_]. Additional notation used to formulate and evaluate the model is summarized in Table 2.

**Table 2.** Summary of notation.

| Notation | Definition |
| --- | --- |
| $n$ | Number of samples |
| $d$ | Dimension of a single sample |
| $m$ | Number of pure breed classes |
| $\theta$ | Purity threshold |
| $X$ | SNP or PC vector of $n$ samples with $\dim(X) = (n, d)$ |
| $X_i$ | SNP or PC vector of a single sample with $\dim(X_i) = (1, d)$ |
| $Y$ | True labels for $n$ samples with $\dim(Y) = (n, m)$ |
| $Y_i$ | True label of a single sample with $\dim(Y_i) = (1, m)$ |
| $\hat{Y}$ | The raw prediction of $n$ samples with $\dim(\hat{Y}) = (n, m)$ |
| $\hat{Y}_i$ | The raw prediction a single sample with $\dim(\hat{Y}_i) = (1, m)$ |
| $\hat{Y}'$ | The transformed prediction of $n$ samples with $\dim(\hat{Y}') = (n, m)$ |
| $\hat{Y}'_i$ | The transformed prediction of a single sample with $\dim(\hat{Y}'_i) = (1, m)$ |
| $I_{PC\_RF_i}$ | PC importance vector of a random forest $\dim(I_{PC\_RF_i}) = (1, d)$ |
| $l$ | Number of SNPs; we also use $l_t$ and $l_b$ to denote the number of SNPs in the top and bottom of the distribution of importance scores |
| $W$ | PCA mapping matrix with $\dim(W) = (d, l)$ |
| $k$ | Window size of SNPs with the highest importance score |
| $j$ | Window size of SNPs with the lowest importance score |

To convert *Ŷ* into a final breed prediction (*Ŷ*′), we applied a post-processing step based on a purity threshold parameter, *θ*, which determines whether an individual is classified as a purebred or mixed-breed dog based on the distribution of predicted breed membership scores. Specifically, the largest element of *Ŷ* is compared with *θ* to determine whether the dog is purebred or mixed-breed (see Methods). The raw prediction vector is then transformed into the corresponding purebred or mixed-breed label format. In this way, the model first produces a continuous breed-membership representation, followed by a discrete prediction that can accommodate both purebred and mixed-breed dogs.

### Evaluation metrics

To evaluate the performance of our framework and compare it with other approaches, we defined two complimentary statistics. Let *INZ*() denote the set of **I**ndices corresponding to **N**on-**Z**ero entries in a label vector. A prediction was considered strictly correct when *INZ*(*Ŷ*_*i*_′) = *INZ*(*Y_i_*), where *Ŷ*_*i*_′ is the final predicted label vector for sample *i* and *Y_i_* is the corresponding owner-reported label vector. A prediction was considered loosely correct when *INZ*(*Ŷ*_*i*_′) ∩ *INZ*(*Y_i_*) ≠ ∅. These summary measures are referred to as strict accuracy and loose accuracy, respectively. In words, strict accuracy requires an exact match between the predicted and reported breed labels, meaning that all breed components of a dog must be correctly identified with no additional or missing breeds. In contrast, loose accuracy is satisfied if the prediction captures at least one of the reported breed components. For example, if a dog is reported as a German Shepherd Dog/Golden Retriever mix, a prediction of German Shepherd Dog alone would be counted as loosely correct but not strictly correct. Because the reference labels were constructed from owner-reported breed information, they should not be regarded as the ground truth, particularly for mixed-breed dogs^22,23^; therefore, the accuracies reported below are conservative.

### Assessing model performance in the Dog Aging Project

In the training set, we evaluated the strict and loose accuracy while varying the purity threshold between 0 < 𝜃 < 1, and found 𝜃 = 0.70 had the highest loose accuracy and strict accuracies of 100% and 99.1%, respectively (Figure 2b). We used this threshold throughout the inference pipeline, and when applied to the independent test set, the model achieved a loose accuracy of 91.7% and a strict accuracy of 58.7%. To visualize model outputs, we constructed a prediction map from the final predicted labels. For clarity, we show a subset of breeds (*m* = 10) in Figure 2c, while the full 100 × 100 prediction map is provided in Figure S1. Because predicted labels can represent breed combinations that do not exactly match the true labels, the same label set is displayed on both axes so that the main diagonal denotes correct predictions.

The prediction map is divided into four quadrants based on whether the true and predicted label correspond to purebred or mixed-breed classes, analogous to a binary confusion matrix.

Quadrant I contains cases where the model predicts purebred and the true label is also purebred. By definition, loose and strict accuracy are identical for predictions in quadrant I. Quadrant II corresponds to cases where the true label is purebred but the predicted label is mixed-breed.

Similarly, quadrant III represents cases where the true label is mixed-breed but the predicted label is purebred. Predictions in quadrants II and III necessarily have a strict accuracy of 0%. Finally, quadrant IV indicates cases where the true and predicted labels are mixed-breed. A summary of the prediction map for 10 breeds is shown in Figure 2c, and Figure 2d summarizes the prediction map of the full dataset (*m* = 100; Figure S1).

Across the full test set (n = 1,979), our model generates 165 loose-wrong predictions. Among these, 73 errors (44.2%) involved an “Unknown” class in either the owner-reported true label or the predicted label, and 45 errors (27.3%) involved a predicted breed that was closely related to or phenotypically similar to the reported breed, such as Miniature American Shepherd and Australian Shepherd, Poodle and Poodle (Toy), Belgian Malinois, Belgian Tervuren and German Shepherd Dog, etc. Overall, 106 errors (64.2%) fell into at least one of these two categories indicating that a substantial proportion of nominal errors likely arose from label noise rather than a failure of model to capture meaningful breed signals. Because all breed assignments were owner reported, the model was likely not trained to its full capability, and the evaluation likely contains many false negatives in which biologically plausible, or effectively correct, predictions were scored as incorrect because of potentially incomplete or inaccurate reported labels.

The second major reason for prediction error is the strong class imbalance in the full dataset, which leaves many breed classes with very little training data. The test set contains 499 breed classes, including 99 purebred classes and 400 mixed-breed classes. Mixed-breed classes were generally represented by fewer samples than purebred classes (Figure 2e). Notably, 283 classes were represented by a single test sample, all of which were mixed-breed classes. By contrast, only 19 classes contained at least 15 test samples (Figure 2e, right), and among these, only four were mixed-breed classes. Two purebred classes, Labrador Retriever (n = 128) and Golden Retriever (n = 125), were especially well represented. Consistent with this skewed distribution, the rarest classes contributed disproportionately to the observed errors (Table 3): test-set classes with only 1 test sample accounted for 239 of 817 strict wrong predictions (29.3%) and 72 of 165 loose wrong predictions (43.6%). More broadly, classes with 5 or fewer test samples comprised 700 of 1,979 test samples, yet contributed 501 strict errors (61.3%) and 133 loose errors (80.6%), whereas classes with more than 15 test samples accounted for only 106 strict (13.0%) and 11 loose errors (6.7%). Thus, the poor performance of rare classes is expected because they are underrepresented not only at evaluation but also during model training, leaving the model with insufficient examples to learn robust breed-specific patterns. Detailed prediction results for each breed class are provided in Table S2. Together, these observations motivated our subsequent focus on breeds with at least 15 test samples, where model performance is less constrained by sample scarcity and downstream genetic analyses are more likely to yield stable and biologically interpretable results.

**Table 3.**
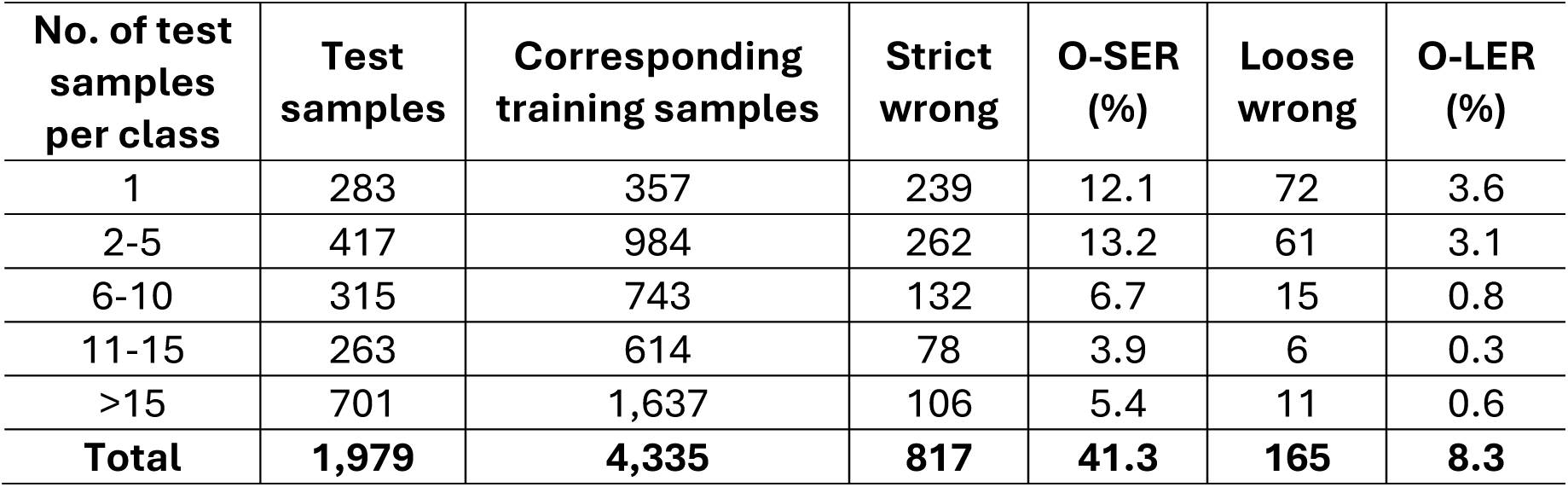
Distribution of strict and loose prediction errors by the number of samples per class in the test set. The corresponding training samples column denotes the number of training samples belonging to the same classes represented in each test-set stratum. These values do not sum to the size of full training-set size (n = 4,593) because 234 mixed-breed classes (n = 258) were exclusive to the training set. O-SER and O-LER denote the contribution to the Overall Strict and Loose Error Rates, respectively.

### Our simple model outperforms ADMIXTURE-based predictions

We next compared our framework with an ADMIXTURE-based benchmark model^13^, which is a popular approach for inferring breed labels. Ancestry estimates from ADMIXTURE were obtained for 1,974 of the 1,979 test samples and converted into the same 100-dimensional label format used in our framework by retaining the two highest-proportion ancestry components and all remaining breed entries were set to zero. The purity threshold, optimized on the training set, was 0.92, yielding the loose accuracy of 87.8% and strict accuracy of 55.7%. Therefore, our simple machine learning framework improved prediction performance by 3.9% in loose accuracy and 3.0% in strict accuracy. Although the improvement is modest, it is notable given ADMIXTURE’s strong performance and the inherent difficulty of distinguishing closely related and admixed breeds. Additionally, these results demonstrate that a simple, interpretable machine learning framework can match and modestly outperform established population-genetic approaches.

### Developing a SNP importance score

A strong motivation for using random forests as the core predictive model is that they provide a natural framework for interpretation, allowing model predictions to be related back to individual genetic variants. To quantify the contribution of each input feature (SNP), we first computed, for each breed-specific random forest, the importance of the 100 principal components (PCs) based on the mean decrease in impurity aggregated across decision trees. This yielded a PC importance vector for breed 𝑖, denoted 𝐼_𝑃𝐶_𝑅𝐹𝑖_, of dimension (1, 100). To account for the unequal contributions of different PCs to the encoded genotype space, we then weighed this vector by the explained variance ratio of the PCA model, resulting in a variance-adjusted PC importance vector, denoted *Ī_PC_RF_i__*.

Next, we projected PC-level importance back to the SNP level using the PCA loading matrix W, which has dimension (100, 54143) and specifies how each of the 54,143 SNPs contributes to each principal component. We therefore defined the SNP importance score for breed *I_SNP_Breed_i__* = *Ī_PC_RF_i__W*, which yields a (1, 54143) vector assigning each SNP a non-negative importance score that reflects its contribution to breed prediction through the learned low-dimensional representation.

### Selection of well-represented breeds for robust downstream analyses

To enable robust downstream genetic analyses, we focused on a subset of 14 purebreds (Figure 3a) with at least 15 individuals in the test set (𝑛 ≥ 15). Restricting the full model (Figure S1) to these classes demonstrated near-perfect predictive performance across 567 purebred test samples, with a loose accuracy of 99.5% (3 incorrect predictions) and a strict accuracy of 91.7% (47 incorrect predictions). Based on this result, we constructed a reduced dataset comprising *m* = 15 classes (14 breeds and an “unknown” class) and n = 2,772 samples. This well-represented subset was used to evaluate predictive performance and to investigate the phenotypic insights provided by SNP importance scores.

**Figure 3.**
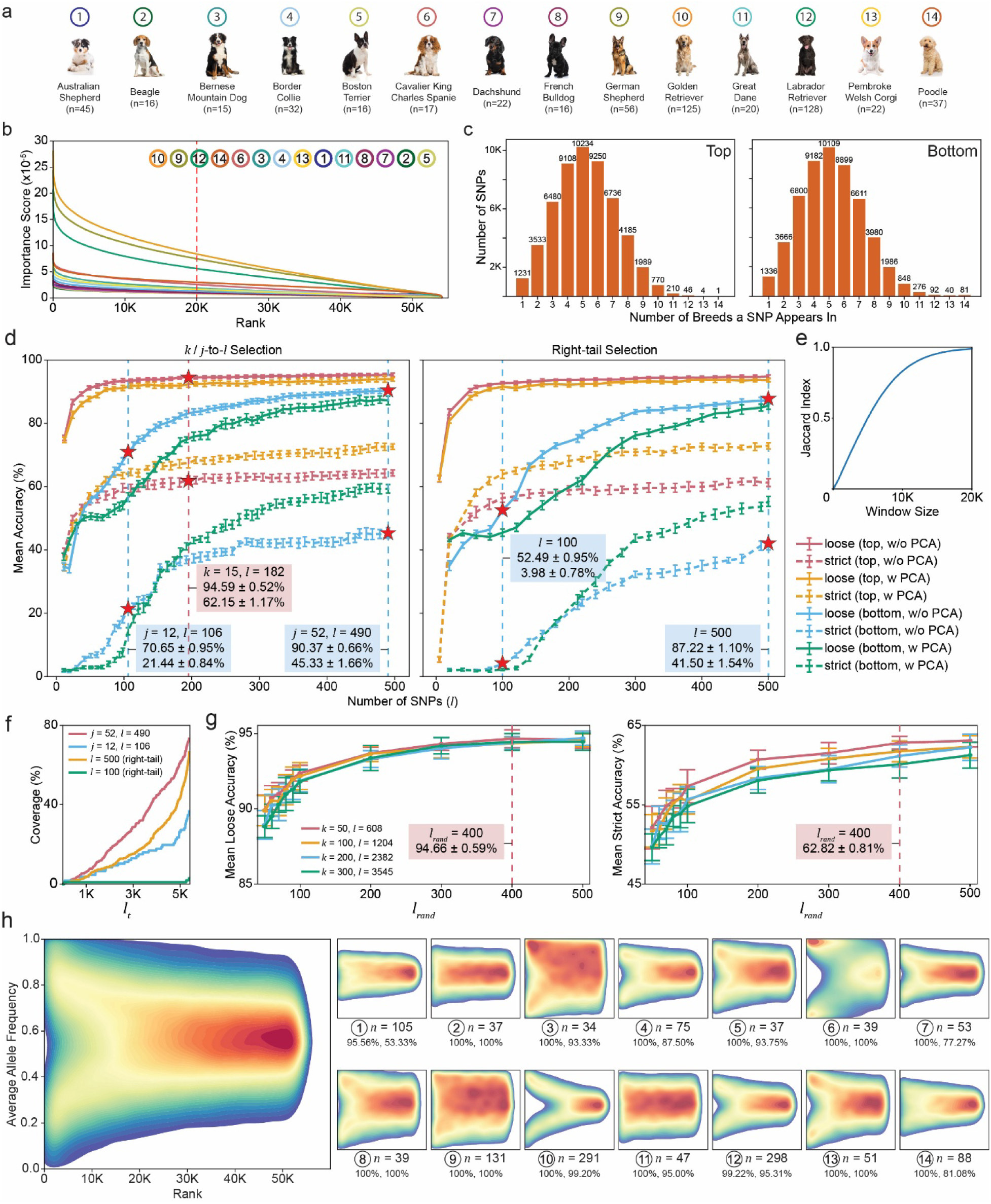
Reduced-SNP prediction and interpretation of informative loci in 14 well-represented breeds. (a) The 14 purebred classes with at least 15 test samples. Values in parentheses indicate sample sizes of the breed in the test set. (b) Breed-specific SNP importance scores ranked in descending order. (c) Distribution of the number of breeds in which a SNP appears within the top ranking window (left, k = 20,000) or bottom ranking window (right, j = 20,000). (d) Mean loose and strict accuracy for reduced SNP panels with and without PCA, using top-ranked and bottom-ranked SNPs under different selection strategies. (e) Jaccard overlap between the unions of top-k and bottom-j SNP sets, with k = j. (f) Coverage of representative bottom-ranked SNP panels by the union of top-k SNP sets. (g) Mean loose accuracy (left) and mean strict accuracy (right) for randomly sampled subsets of top-ranked SNPs. (h) Heatmap SNP-importance rank versus average allele frequency across all breeds (left) and within each breed (right). Color denotes the 2D kernel density estimate of SNP distribution in the space defined by SNP importance rank and average allele frequency, where warmer colors indicate higher density and cooler colors indicate lower density.

### Leveraging SNP importance scores to create small panels of informative variants

We next asked how small a reduced SNP panel could be while still retaining substantial predictive power. The SNP importance scores provide a natural framework for identifying a small panel of informative SNPs. Because SNP importance scores were computed separately for each breed, the informativeness of a given SNP is inherently breed-specific. This overall pattern is shown in Figure 3b, which displays the 54,143 SNPs ranked by importance score for each of the 14 well-represented breeds. As one illustrative example, chr17:63149205:T:C ranked 53,345th of 54,143 in Cavalier King Charles Spaniel but ranked first in both German Shepherd Dogs and Golden Retrievers. Notably, the average allele frequencies of this SNP in the latter two breeds were 0.04 and 0.96, respectively, indicating that the same variant can be highly informative in multiple breeds while still strongly distinguishing between them.

We first considered a breed-specific SNP selection strategy in which, for each breed, the top *k* SNPs ranked by importance were selected and then combined across all breeds by taking their union. Because many SNPs were shared among breeds, the resulting set contained *l* unique SNPs (with 𝑙 ≤ 𝑚 × 𝑘), which we refer to as *k*-to-*l* selection (*m* = 14 as we are focusing on well-represented breeds). This strategy prioritizes SNPs that are highly informative within at least one breed, while allowing overlaps across breeds to reduce redundancy.

We also considered a complementary strategy to identify a small panel of informative SNPs that prioritize highly ranked variants shared across multiple breeds. Specifically, for each SNP, we counted the number of breeds in which it appeared within a predefined importance-ranking window (i.e., the top *k* SNPs per breed). This yielded a distribution reflecting how broadly each SNP contributed to breed prediction (Figure 3c). SNP panels were then constructed by selecting variants from the right tail of this distribution—that is, SNPs that were highly ranked in many breeds. This strategy favors SNPs capturing genetic variation that is informative about ancestry across multiple breeds rather than variants that are highly specific to a single breed. We refer to this procedure as right-tail selection. These two strategies capture complementary aspects of breed-associated genetic variation: *k*-to-*l* selection prioritizes SNPs that are highly informative within individual breeds, whereas right-tail selection enriches for variants that are consistently informative across multiple breeds.

### Accurately predicting breed labels from several hundred informative SNPs

To evaluate the predictive accuracy of these reduced SNP panels, we increased the testing proportion to 0.5, leaving 1,386 samples for training in the *m* = 15 dataset. Because the resulting training set was relatively small, we limited the panel size to 𝑙 ≤ 500. For *k*-to-*l* selection, we evaluated 1 ≤ 𝑘 ≤ 41, which yielded 𝑙 values ranging from 12 to 497. For right-tail selection, we evaluated 𝑙 from 0 to 500 in increments of 20, replacing floor-multiple values with exact bracket sizes where appropriate. These additional values were 5, 51, and 261. As a separate input representation, we also applied PCA to each reduced SNP panel, retaining the number of principal components required to explain 95% of the variance.

For each SNP panel, we trained the same prediction framework as described above, namely a collection of *m* breed-specific random forests with 100 decision trees per forest. The dataset was randomly partitioned 20 times into equal training and test sets, and for each partition, we recorded the mean and standard deviation of loose and strict accuracy across four settings: *k*-to-*l* selection with and without PCA, and right-tail selection with and without PCA. In total, 1,680 models were trained under *k*-to-*l* selection and 1,040 under right-tail selection (Figure 3d).

Across all four settings, performance increased rapidly and then plateaued as the number of selected SNPs increased. Without PCA, *k*-to-*l* and right-tail selection reached approximate loose accuracies of 95.3% and 94.8%, and strict accuracies of 64.3% and 61.6%, respectively.

Applying PCA led to a modest decrease in loose accuracy of 93.9% and 93.6%, but the strict accuracy substantially improved to 72.6% and 72.9%, respectively. Thus, PCA narrowed the gap between loose and strict accuracy. Overall, *k*-to-*l* selection performed slightly better than right-tail selection, suggesting that SNPs ranked highly within a subset of breeds are somewhat more informative than SNPs shared more broadly but ranked lower within a breed.

Notably, 99% of the plateau effect was achieved with only 157 and 305 SNPs for *k*-to-*l* selection without and with PCA, and 240 and 261 SNPs for right-tail selection without and with PCA, respectively. These results indicate that, in this setting, approximately 150 to 300 informative SNPs are sufficient for a near-maximal prediction performance.

Despite improved performance with top-ranked SNPs, the corresponding SNP sets showed substantial overlap at larger window sizes. For example, the union of the top 20,000 SNPs across all 14 breeds contained 53,777 SNPs, corresponding to 99.3% of the full set of 54,143 SNPs.

Similarly, the union of the bottom 20,000 SNPs contained 53,906 SNPs (99.6% of all SNPs), and the overlap between the top- and bottom-ranked unions was extensive (Jaccard index = 98.9%; Figure 3e).

To determine whether the performance of top-ranked SNP panels simply reflected this broad overlap, we repeated the analysis using bottom-ranked SNPs as a control. Analogous to the top-ranked setting, we defined *j-to-l* selection as a strategy in which, for each breed, the top *j* SNPs (here drawn from the bottom-ranked set) were selected and then combined across breeds, yielding *l* unique SNPs after taking the union. We also considered a corresponding right-tail-bottom selection procedure based on SNPs that were consistently ranked among the least informative across breeds.

For *j-to-l* selection, we evaluated 1 ≤ 𝑗 ≤ 52, which produced *l* values ranging from 11 to 490. For right-tail-bottom selection, we evaluated *l* from 20 to 500 in increments of 20, replacing floor-multiple values with exact bracket sizes where necessary (81, 121, 213, and 489). Model fitting and dataset partitioning were performed as described above.

Prediction using bottom-ranked SNP panels was consistently worse than prediction using top-ranked panels (Figure 3d). However, performance improved as panel size increased, consistent with the rapid growth in overlap between the top-ranked and bottom-ranked unions shown in Figure 3e. To examine this more directly, we measured how strongly representative bottom-ranked SNP panels overlapped with top-ranked SNP sets as the top-ranked window size increased. Specifically, for a given bottom-ranked SNP panel 𝑆_𝑏_, we denote its size by 𝑙_𝑏_ = |𝑆_𝑏_|. For each value of 𝑘, we then constructed 𝑆_𝑡_ as the union of the top-𝑘 SNP sets across all 14 breeds and denote its size by 𝑙*_t_* = |𝑆*_t_*|. Coverage was defined as 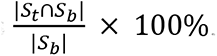. As 𝑘 increased from 10 to 20,000, 𝑙_𝑡_ increased accordingly, reaching 53,777 when 𝑘 = 20,000. The better-performing bottom-ranked panels, 𝑗 = 52 with 𝑙_𝑏_ = 490 and right-tail-bottom selection with 𝑙_𝑏_ = 500, shared 361 of 490 SNPs (73.67%) and 331 of 500 SNPs (66.20%), respectively, with the corresponding top-ranked unions. By contrast, the poorer-performing panels, 𝑗 = 12 with 𝑙_𝑏_ = 106 and right-tail-bottom selection with 𝑙_𝑏_ = 100, shared only 39 of 106 SNPs (36.79%) and 3 of 100 SNPs (3.00%), respectively (Figure 3f). Together, these control analyses support the conclusion that high-ranking SNPs are genuinely enriched for predictive information, rather than benefiting only from broad set overlap. Detailed results are provided in Tables S3-S6.

### Toward portable SNP panels for breed prediction

A practical limitation of the full model is that it requires the exact same set of 54,143 SNPs used during training, so that the learned PCA transformation and random forest models can be applied without modification. This requirement is restrictive in practice, particularly when analyzing data generated from different genotyping platforms or sequencing pipelines, where SNP overlap may be incomplete.

To explore a more portable alternative, we evaluated whether accurate prediction could be achieved using randomly sampled subsets of informative SNPs. Using the reduced dataset described above (*m* = 15, *n* = 2,772), we fixed the training–testing split at 1:1 and constructed four pools of top-ranked SNPs using *k*-to-*l* selection with 𝑘 ∈ {50, 100, 200, 300}, yielding 𝑙 = 608, 1204, 2382, and 3545 SNPs, respectively. From each pool, we randomly sampled panels of size 𝑙_𝑟𝑎𝑛𝑑_ ∈ {50, 60, 70, 80, 90, 100, 200, 300, 400, 500} SNPs, with 20 independent replicates per combination. Because these panels contained at most 500 SNPs, PCA was not applied in this analysis. Model fitting and evaluation were otherwise performed as described above and mean loose accuracy across conditions is shown in Figure 3g.

Although the most favorable scenario—namely, that very small random panels drawn from large SNP pools would retain near-maximal performance—was not observed, several informative trends emerged. First, smaller *k* values consistently yielded better performance, indicating that SNP pools restricted to the highest-ranked variants are more enriched for breed-discriminating signal. This result provides additional empirical support for the SNP importance score as a meaningful measure of predictive information. Second, prediction accuracy increased with panel size 𝑙_𝑟𝑎𝑛𝑑_, with performance approaching a plateau for panels of a few hundred SNPs. These results indicate that relatively small sets of informative SNPs, even when selected at random, can capture much of the predictive signal in the data.

Comparison with importance-prioritized panels (Figure 3d) highlights a trade-off between portability and efficiency. Panels constructed by explicit prioritization achieve high accuracy with relatively few SNPs, whereas randomly sampled panels require larger sizes to achieve comparable performance. For example, selecting the top 15 SNPs per breed yielded a panel of 182 SNPs with a loose accuracy of 94.6 ± 0.52% and strict accuracy of 62.2 ± 1.17%. In contrast, a randomly sampled panel required 400 SNPs drawn from a pool of 608 top-ranked variants to achieve similar performance (loose accuracy 94.7 ± 0.59%, strict accuracy 62.8 ± 0.81%). Together, these results demonstrate that portable SNP panels can achieve strong predictive performance, albeit at the cost of reduced efficiency relative to importance-ranked panels.

### Informative SNPs identify known and novel loci contributing to hallmark breed phenotypes

Gene coordinates and annotations were obtained from the Dog10K annotation resource^24,25^. SNPs were assigned to genes when their genomic coordinates fell within the annotated span of a gene. We first examined pairwise overlap among the top 20,000 SNPs for the 14 breeds. As shown in Figure S2, most breed pairs share approximately 7,000 to 7,700 SNPs, corresponding to Jaccard indices of about 0.22-0.25. By contrast, two pairs stood out clearly from the background distribution: German Shepherd Dog and Golden Retriever shared 19,747 SNPs (Jaccard index = 0.98), and Beagle and Dachshund shared 19,190 SNPs (Jaccard index = 0.92). These exceptional overlaps suggest that breed prediction for these pairs draws on highly similar sets of informative loci.

We next focused on a more stringent subset comprising the top 3,000 shared SNPs within each pair. Among these shared high-ranking SNPs, several mapped to genes previously associated with canine phenotypes (Table 4). For the German Shepherd Dog-Golden Retriever pair, these included *HMGA2* and *ADAMTS9*, which have been linked to body size and weight^26,27^, *PTPRD*, which has been implicated in hip dysplasia^28,29^, and *RSPO2*, *FOXI3*, and *CA10*, which are associated with coat or hair-related phenotypes^26,30,31^. An additional shared locus mapped to *EPHA5*, which has previously been associated with sheepdog-related traits, such as herding behavior^11^. For the Beagle-Dachshund pair, shared high-ranking SNPs mapped to genes associated with curl tail (*CHSY3*), facial morphology (*ZFHX3*), pigmentation (*ASIP*), body weight (*R3HDM1*) and hip dysplasia (*PTPRD*), together with *CA10*, which has been linked to hairlessness^26,28,29,31^. Together, these results indicate that the shared predictive signal between these breed pairs is enriched in loci related to morphology, skeletal traits, and other breed-associated phenotypes.

**Table 4.**
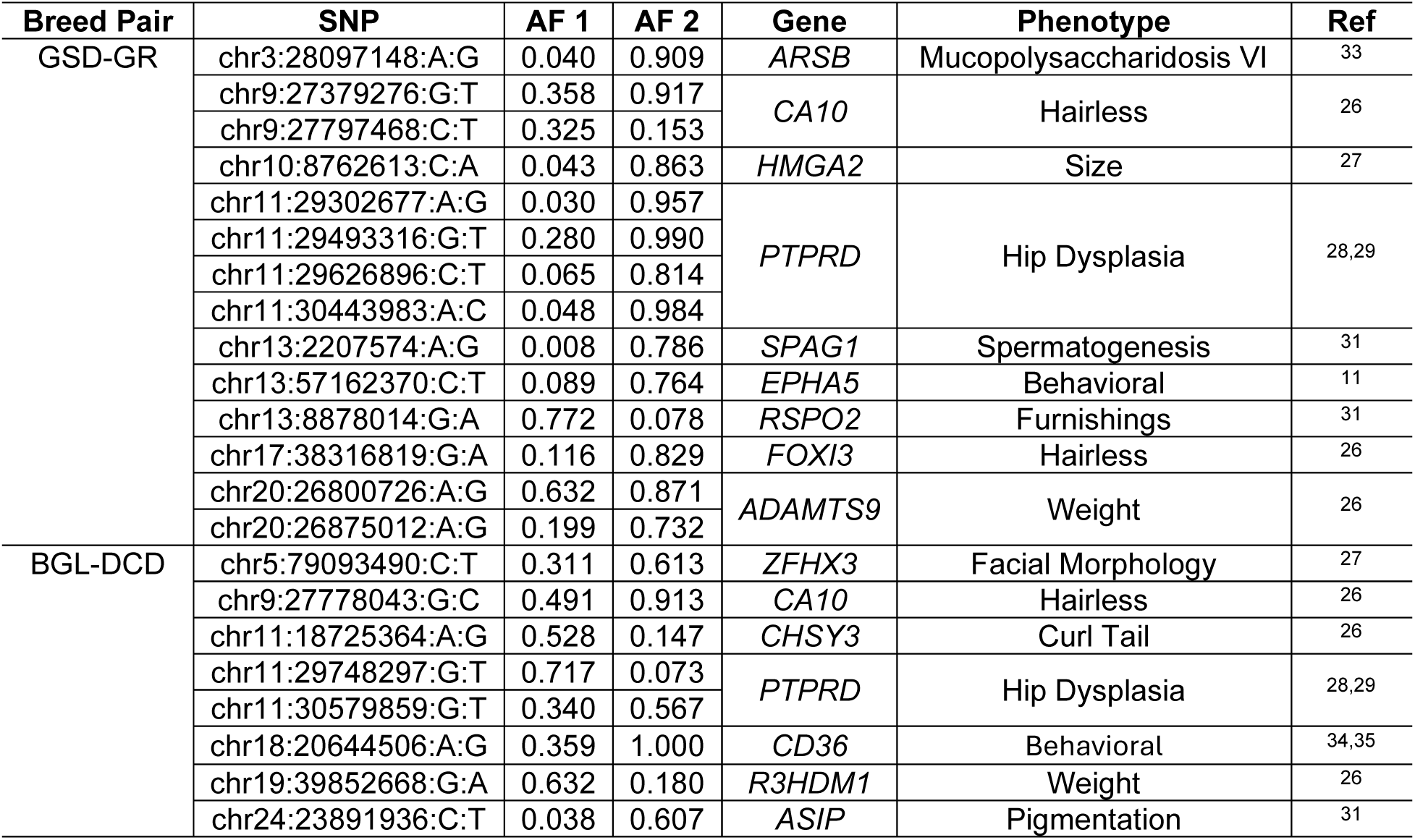
Shared high-ranking SNPs in two breed pairs that map to genes associated with phenotypes. The first breed pair is the German Shepherd Dog (GSD, breed 1) and Golden Retriever (GR, breed 2) and the second breed pair is the Beagle (BGL, breed 1) and Dachshund (DCD, breed 2). AF1 and AF2 denote allele frequencies in breed 1 and breed 2, respectively.

Both breed pairs also contained highly ranked shared SNPs without prior phenotype annotation (Table 5). These variants ranked among the top contributors to breed prediction in both breeds within a pair yet were either not mapped to an annotated gene or lacked a previously reported canine phenotype association. Notably, such variants remained highly informative despite the absence of established functional annotation, highlighting the ability of the framework to identify candidate loci that would not be prioritized by prediction accuracy alone. This feature further illustrates the interpretability of the model, as it recovers both biologically established loci and putatively novel breed-associated signals.

**Table 5.**
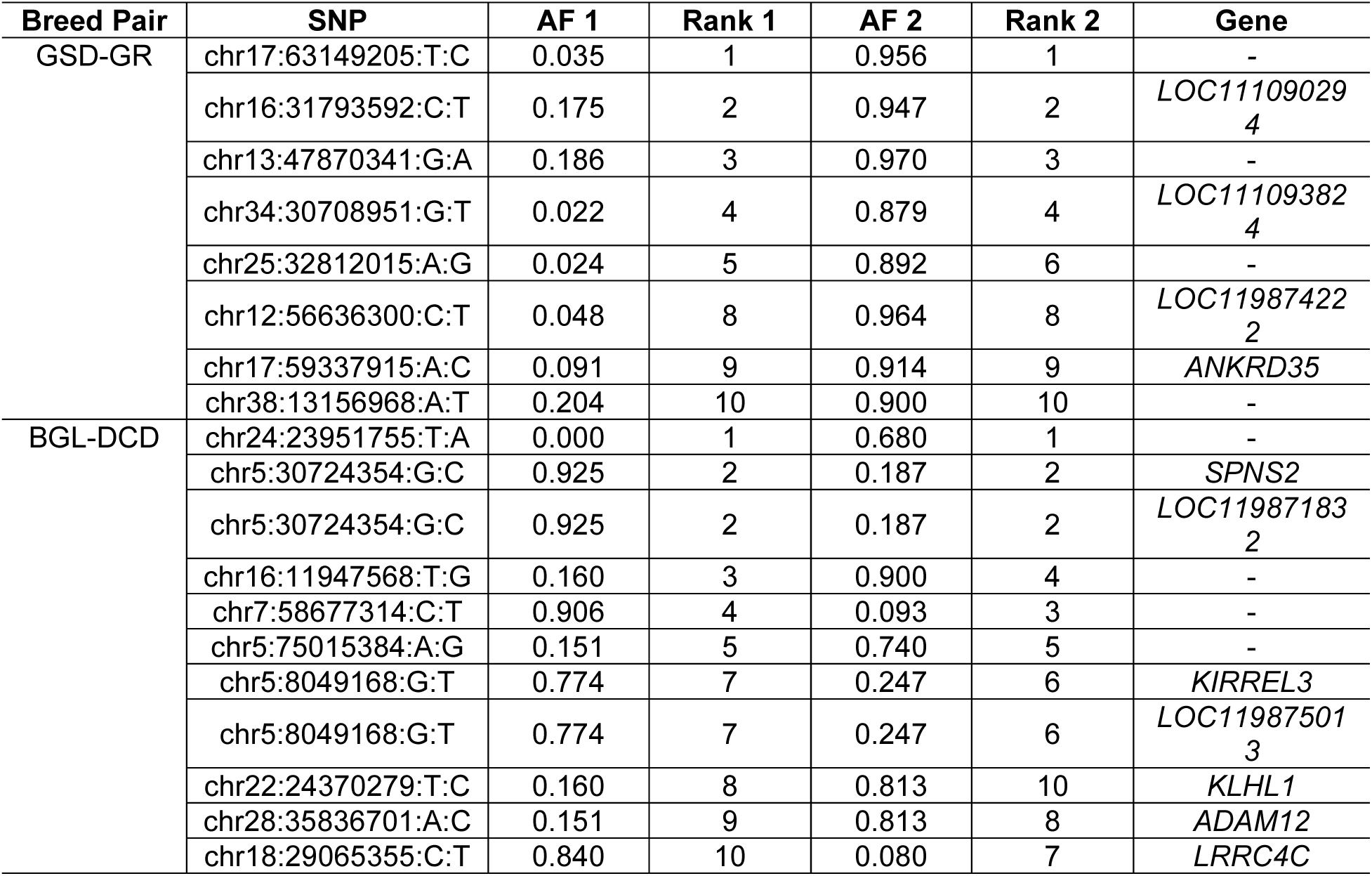
Shared high-ranking SNPs in two breed pairs that map to loci with unknown phenotypic consequences. The notation is the same as in Table 2. Rank1 and Rank2 denote the within-breed importance ranks of the SNPs for breed 1 and breed 2, respectively.

Extending this analysis across all 14 breeds and restricting attention to SNPs within the top 1,000 ranking window, we identified 35 SNPs mapping to 20 genes previously associated with reported canine phenotypes (Table 6). Several recurrent signals were biologically plausible in the context of breed differentiation, including *HMGA2*, *GHR*, *MED13L*, *RNFT2*, and *HNF4G*, which have been linked to body size, height, or bulk^26,27,31^; *RSPO2* and *ASIP*, which are associated with coat furnishings and pigmentation^30,31^; and *CHSY3* and *PTPRD*, which are linked to tail morphology and hip dysplasia^26,28,29^. Beyond these morphology- and skeletal trait-associated loci, *EFNA5* has previously been associated with sheepdog-related signals^11^, whereas loci such as *FTO*, *FGR*, *SPAG1,* and *CD36* are less directly connected to canonical breed-defining morphology. These latter associations may reflect pleiotropy, linkage to nearby causal variants or broader haplotypic signatures characteristic of particular breeds. Overall, the enrichment of highly ranked SNPs near genes with known effects on morphology, pigmentation, skeletal traits and breed-associated biology supports the biological interpretability of the SNP importance score.

**Table 6.**
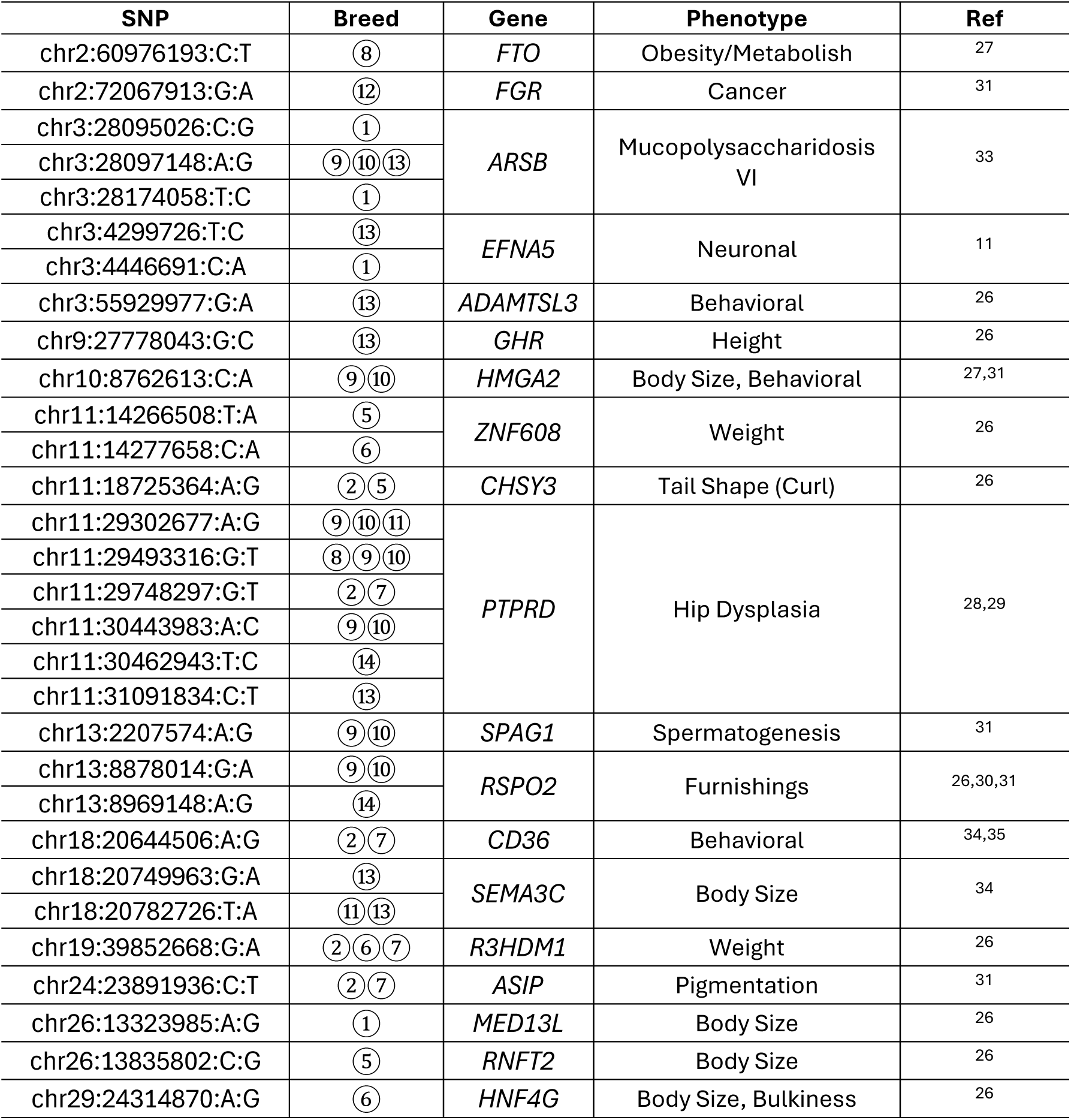
SNPs within the top 1,000 importance-ranked variants that map to genes with known phenotypic associations. Breed indices correspond to the 14 breeds shown in Figure 3a.

## Discussion

We developed an interpretable framework for inferring dog breed labels from genome-wide SNP data and used it to investigate the structure of breed-associated genetic variation. Using the DAP dataset^20^, we found that accurate breed prediction can be achieved using a highly compressed genomic representation, with predictive performance remaining stable despite substantial reductions in both SNP dimension and feature space. Indeed, a striking finding of this study is that only a few hundred informative SNPs are sufficient to achieve near-maximal predictive performance. Reduced-panel analyses demonstrated that carefully selected SNP sets retain most of the predictive signal present in the full model, while even randomly sampled subsets of informative variants can achieve strong performance when the panel size is sufficiently large. These results indicate that breed-discriminating information is both widely dispersed and highly redundant across the canine genome, reflecting the effects of artificial selection, inbreeding, and genetic drift during breed formation^1–3^.

We evaluated a range of parameter values to optimize our model’s performance (Table 1).

The final encoding parameters were selected to balance compactness, robustness, and predictive performance. Notably, across the parameter settings examined, model accuracy remained stable despite large changes in SNP and PCA dimensions (Table 1). We therefore prioritized a configuration that preserved the strong predictive performance of our model while keeping both the SNP representation and the PCA embedding relatively small. Although the most aggressive setting tested, MAF < 0.50, resulted in the smallest SNP dimension without an obvious loss of accuracy, we regarded this threshold as overly restrictive for the initial whole-genome encoding because it yields a highly constrained feature space dominated by common variants, potentially limiting the robustness and transferability of the encoding across datasets and downstream analyses.

The SNP importance scores derived from our machine learning framework model provide additional insight into the genetic architecture underlying breed differentiation. Highly ranked SNPs tend to exhibit more extreme allele frequencies, consistent with the expectation that loci distinguishing breeds are often driven toward fixation within breeds while remaining differentiated across populations. However, given the unique history of domesticated dogs and the strong correlation among variants^10^, most informative SNPs are not causally related to breed-defining phenotypes but instead reflect LD with targets of artificial selection (or highly differentiated variants resulting from intense genetic drift following population bottlenecks^1,3^).

Nevertheless, the strong association between highly informative SNPs and genes associated with known canine traits supports the biological relevance of the learned signals and highlights the potential of the framework for generating hypotheses.

Our analyses also reveal a practical trade-off between efficiency and portability in reduced-SNP panels. Panels constructed by explicitly prioritizing informative variants achieve high accuracy with relatively few SNPs but depend on the specific variants selected and may thus not transfer readily across datasets. In contrast, panels constructed by random sampling from informative SNP pools are more flexible and robust to the specific set of variants chosen but require more SNPs to achieve comparable performance. This trade-off has important implications for the design of portable breed-prediction assays and suggests that hybrid strategies may offer a useful practical compromise.

Our study has several important limitations. For example, the current framework adopts relatively simple representations of mixed-breed labels and classification thresholds, which could be refined in future work to support more quantitative modeling of ancestry composition.

Additionally, we focused on identifying the most dominant components of ancestry in mixed-breed dogs. Although it would be relatively straightforward to extend the modeling framework to allow predicting ancestry from more than two breeds, this is a limitation of our model as currently formulated. If inferences of more minor components of breed ancestry are needed, ADMIXTURE^13^ is likely the preferred methodological framework as, in theory, it can handle the identification of any number of breed classes given a sufficiently comprehensive reference dataset. However, to our knowledge, there have been no systematic studies evaluating the accuracy of breed inference in mixed-breed dogs when there are three or more ancestry components. Thus, this is an important methodological direction for future work. Finally, the reduced SNP panels were derived from the filtered subset of 54,143 variants rather than the full spectrum of canine genetic variation. Therefore, it may be possible to construct better performing minimal SNP panels, although the redundant signal of breed-discrimination we find across the canine genome suggests that any increase in performance from using unfiltered SNPs will be modest.

Despite these limitations, our results demonstrate that accurate and interpretable breed prediction can be achieved using compact genomic representations without relying on strong population-genetic assumptions^14–16^. More broadly, this work highlights the highly structured and compressible nature of canine genetic variation, provides a foundation for portable reduced-SNP prediction panels, and offers a framework for linking predictive models to biologically meaningful genetic features. Our results have potential applications in veterinary genomics, ancestry inference, delineating the genetic architecture of canine phenotypic variation, and the study of genetic diversity in domesticated populations.

## Methods

### Dataset curation and analysis

Low-coverage whole-genome sequencing data were obtained from the 2023 release of the Dog Aging Project^20^, and all project data are housed on the Terra platform at the Broad Institute of MIT and Harvard. Reads were aligned to the UU_Cfam_GSD_1.0 reference genome, and missing genotypes were imputed using GLYMPSE2^32^. Restricting the dataset to biallelic SNPs yielded a final PLINK set containing 7,627 dogs and 29,089,701 SNPs. Breed information for genotyped dogs was provided by DAP participants through survey forms. Each dog was reported as either purebred (n = 4,112) or mixed-breed (n = 3,514). Examples of mixed-breed labels included “German Shepherd Dog / Golden Retriever”, “Dachshund / Unknown”, and “Unknown / Unknown”. During curation, we removed eight samples with inconsistent metadata (Table S7), such as dogs labeled as “Border Collie/Border Collie” but reported as mixed-breed. We also removed one sample (dog_id = 27669) because it caused PLINK to fail during processing.

Finally, we excluded eight samples labeled as village dogs because they are genetically distinct from the purebred populations considered in this study. After data QC, 7,618 samples remained.

For LD pruning, we used 𝑟^2^with a window size of 500 SNPs and a step size of 50 SNPs. To optimize the purity threshold, 𝜃, we performed an exhaustive search over the interval [0, 1] with increments of 0.01, using the training set to compare predicted labels with owner-reported labels. We quantified the resulting loose and strict accuracies across threshold values, which are shown in Figure 2b. We selected 𝜃 using a simple rule: among all threshold values that achieved the maximum loose accuracy, we chose the one with the highest strict accuracy.

### Multi-Output Random Forest Regressor

The prediction model consists of 𝑚 independent random forest regressors, one for each breed class. In practice, this was implemented using MultiOutputRegressor(RandomForestRegressor(n_estimators=100, random_state=42) in scikit-learn. Here, n_estimators = 100 specifies the number of decision trees in each breed-specific random forest, and random_state = 42 was used to improve reproducibility of bootstrap sampling and feature selection during model fitting.

### Visualization of Principal Components

The final SNP encoding compressed 54,143 SNPs into 100 principal components, which together explained 24.63% of the variance in the post-filtering SNP matrix. To visualize the resulting low-dimensional structure, we generated interactive three-dimensional scatter plots of the top three principal components using scatter_3d() in Plotly. These visualizations were produced for five subsets of test samples. For example, pca_breed_plot_pure.html displays all purebred test samples colored by breed. Several breeds, including Labrador Retriever, German Shepherd Dog, Golden Retriever, Flat-Coated Retriever, Newfoundland and Poodle, formed distinct clusters. One Belgian Malinois sample (dog_id = 29084) fell within the German Shepherd Dog cluster, which is not unexpected given the close phenotypic similarity between these breeds. All interactive PCA plots are provided in Supplementary Information (PCA_visualization.zip).

### Source code availability

All source code necessary to install, run, and interpret the breed prediction method described in this manuscript is available at https://github.com/AkeyLab/DAP_breed_prediction.git.

## Acknowledgements

We acknowledge members of the Akey laboratory for constructive feedback on the manuscript, participants of the Dog Aging Project, their dogs, and community veterinarians, as well as all Dog Aging Project Team members (2025) [https://dogagingproject.org/team-acknowledgements] for their important contributions. This work was supported by NIH grant U19 AG057377 (DELP) from the National Institute on Aging, and by additional grants and private donations, including generous support from the Glenn Foundation for Medical Research, the Tiny Foundation Fund at Myriad Canada, the WoodNext Foundation, and the Dog Aging Institute.

## Dog Aging Project Consortium

Joshua M. Akey¹, Rozalyn M. Anderson², Elhanan Borenstein³, Marta G. Castelhano⁴, Amanda E. Coleman⁵, Kate E. Creevy⁶, Matthew D. Dunbar⁷, Virginia R. Fajt⁸, Jessica M. Hoffman⁹, Erica C. Jonlin¹⁰, Matt Kaeberlein¹⁰, Elinor K. Karlsson¹¹, Kathleen F. Kerr¹², Jing Ma¹³, Evan L. MacLean¹⁴, Stephanie McGrath¹⁵, Natasha J. Olby¹⁶, Daniel E.L. Promislow¹⁷, May J. Reed¹⁸, Audrey Ruple¹⁹, Stephen M. Schwartz²⁰, Sandi Shrager²¹, Noah Snyder-Mackler²², M. Katherine Tolbert⁶

1. Lewis-Sigler Institute for Integrative Genomics, Princeton University, Princeton, NJ, USA
2. University of Wisconsin Madison, Madison, WI, USA
3. Department of Clinical Microbiology and Immunology, Gray Faculty of Medical and Health Sciences, Tel Aviv University, Tel Aviv, Israel
4. Cornell Veterinary Biobank, College of Veterinary Medicine, Cornell University, Ithaca,
5. Department of Small Animal Medicine and Surgery, College of Veterinary Medicine, University of Georgia, Athens, GA, USA
6. Department of Small Animal Clinical Sciences, Texas A&M University School of Veterinary Medicine & Biomedical Sciences, College Station, TX, USA
7. Center for Studies in Demography and Ecology, University of Washington, Seattle, WA,USA
8. Department of Veterinary Physiology and Pharmacology, Texas A&M University School of Veterinary Medicine & Biomedical Sciences, College Station, TX, USA
9. Department of Biological Sciences, Augusta University, Augusta, GA, USA
10. Department of Laboratory Medicine and Pathology, University of Washington School of Medicine, Seattle, WA, USA
11. Genomics & Computational Biology, UMass Chan Medical School, Worcester, MA, USA
12. Department of Biostatistics, University of Washington, Seattle, WA, USA
13. Division of Public Health Sciences, Fred Hutchinson Cancer Research Center, Seattle, WA, USA
14. College of Veterinary Medicine, University of Arizona, Tucson, AZ, USA
15. Department of Clinical Sciences, College of Veterinary Medicine and Biomedical Sciences, Colorado State University, Ft. Collins, CO, USA
16. Department of Clinical Sciences, College of Veterinary Medicine, North Carolina State University, Raleigh, NC, USA
17. Jean Mayer USDA Human Nutrition Research Center on Aging at Tufts University, Boston, MA, USA
18. Department of Medicine, Division of Gerontology and Geriatric Medicine, University of Washington School of Medicine, Seattle, WA, USA
19. Department of Population Health Sciences, Virginia-Maryland College of Veterinary Medicine, Virginia Tech, Blacksburg, VA, USA
20. Epidemiology Program, Fred Hutchinson Cancer Center, Seattle, WA, USA
21. Collaborative Health Studies Coordinating Center, Department of Biostatistics, University of Washington, Seattle, WA, USA
22. School of Life Sciences, Arizona State University, Tempe, AZ, USA

