## Supplementary figures and images for "An interpretable machine learning framework for dog breed inference and ancestry decomposition"

### Fig S1

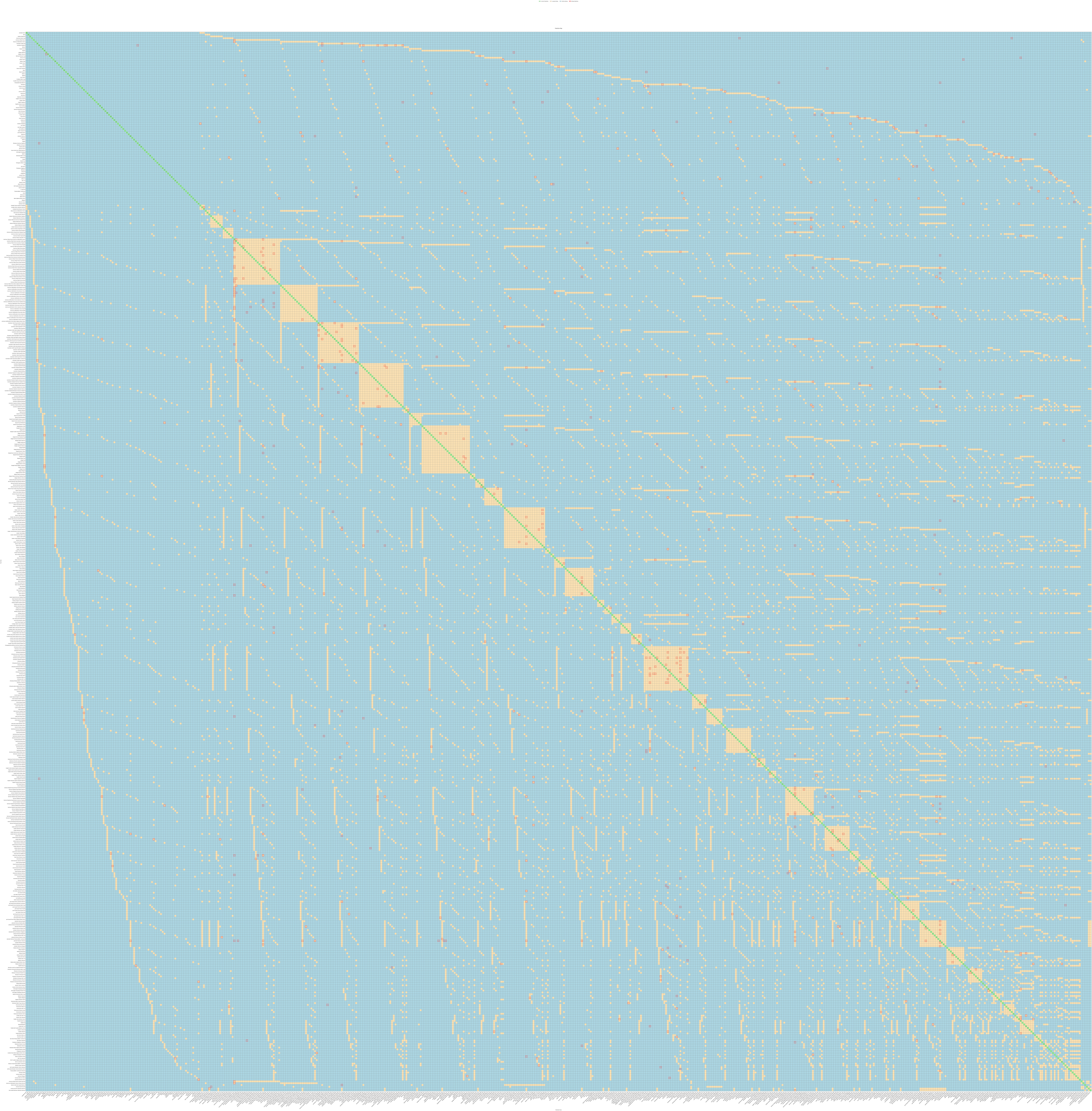

### Fig S2

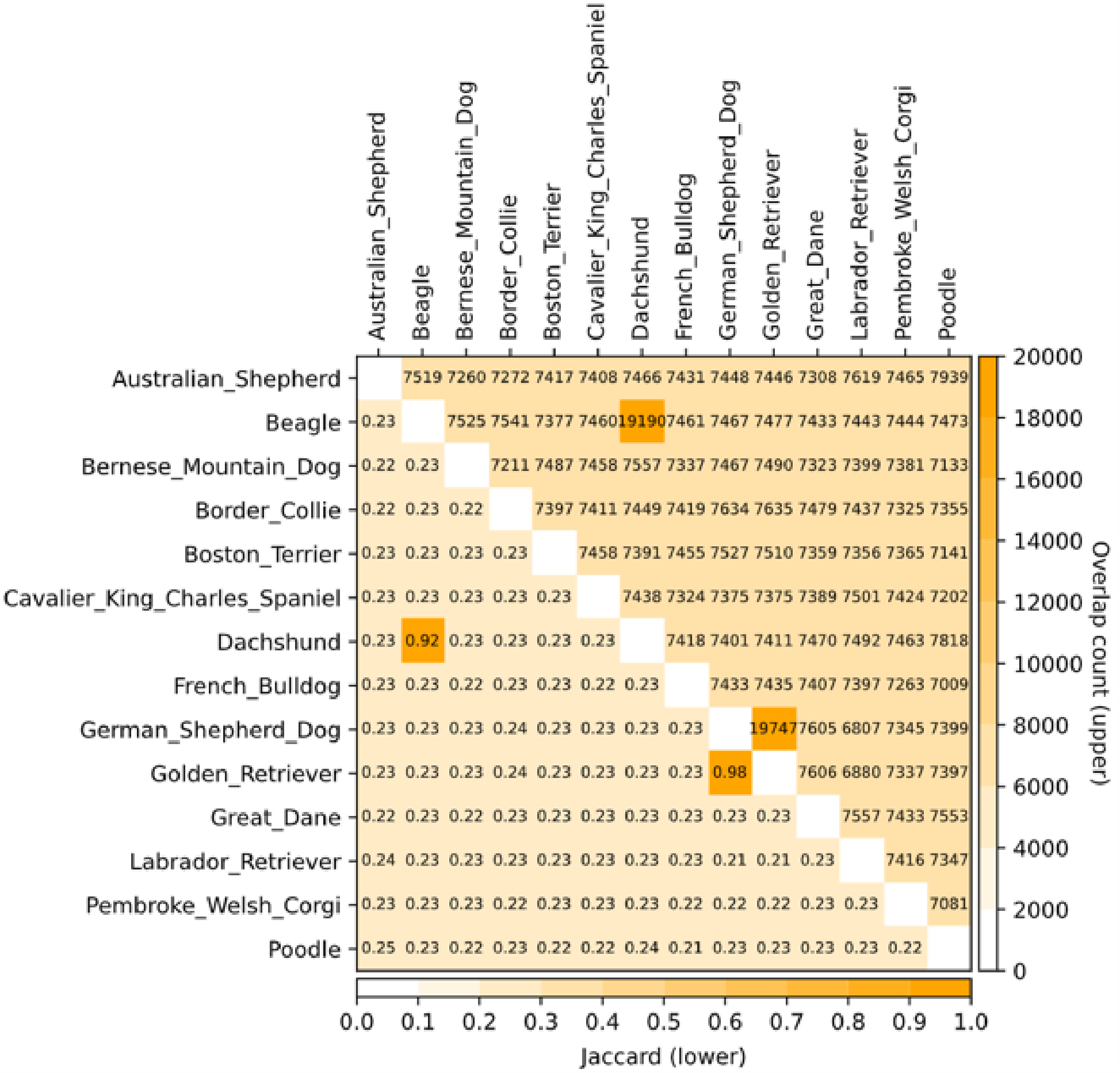
